# Construction and Characterisation of a Structured, Tuneable, and Transparent 3D Culture Platform for Soil Bacteria

**DOI:** 10.1101/2023.06.28.546105

**Authors:** Liam M. Rooney, Lionel X. Dupuy, Paul A. Hoskisson, Gail McConnell

**Author notes:** Department of Conservation of Natural Resources, Neiker, Basque Institute for Agricultural Research and Development, Derio, Spain and Ikerbasque, Basque Foundation for Science, Bilbao, Spain (Lionel X. Dupuy).

## Abstract

2.

We have developed a tuneable workflow for the study of soil microbes in an imitative 3D soil environment that is compatible with routine and advanced optical imaging, is chemically customisable, and is reliably refractive index matched based on the metabolic profile of the study organism. We demonstrate our transparent soil pipeline with two representative soil organisms, *Bacillus subtilis* and *Streptomyces coelicolor*, and visualise their colonisation behaviours using fluorescence microscopy and mesoscopy. This spatially structured, 3D approach to microbial culture has the potential to further study the behaviour of other difficult-to-culture bacteria in conditions matching their native environment and could be expanded to study microbial interactions, such as interaction, competition, and warfare.

**Graphical Abstract:** 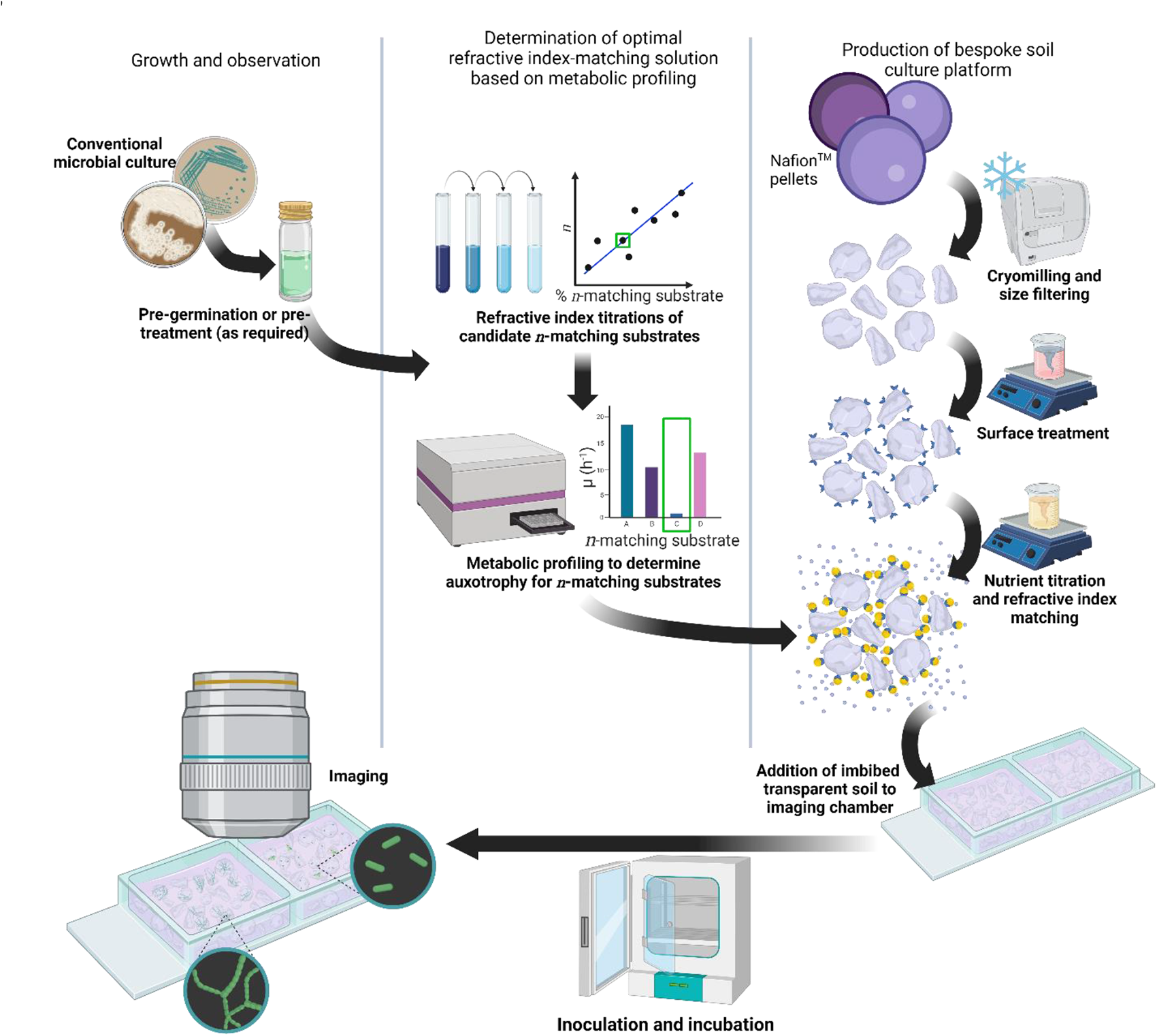

**A step-by-step method for creating a tailored 3D culture medium for study of soil microbes.**

The complete workflow can be split into three parts: Growth and observation, metabolic profiling to provide a stable refractive index matching solution, and production of the 3D soil environment. The 3D culture scaffold was created by cryomilling Nafion™ resin pellets and size filtration. Chemical processing altered the surface chemistry of Nafion™ particles and facilitated nutrient binding by titration of a defined liquid culture medium. Metabolic profiling determined non-metabolisable sugars and provided an inert refractive index matching substrate, which was added to the final nutrient titration. Inoculation and growth of the test strain allowed for downstream assessment of colonisation behaviours and community dynamics *in situ* by, for example, optical microscopy.

## 4. Introduction

Soil is heterogeneous(1) and its complexity presents a challenge for microbial ecologists – how to study the natural behaviours of microorganisms in a representative culture environment, particularly by means of optical microscopy?

Most phenotypic observations of soil bacteria are performed using century-old 2D culture techniques that are unsuitable for *in situ* optical imaging and are neither environmentally nor physiologically representative. The mammalian cell biology field was revolutionised using 3D culture methods over the last 20 years, facilitating discoveries in cell migration and chemotaxis, cell signalling, and tumour development(2–11). However, these 3D culture methods have been poorly adapted for microbiology applications.

*In situ* microscopy of soil microbes have been impeded due to the incompatible material properties of soil. These include compositional variation(1,12–14), the presence of bacteriophage and other competing microbes (which can be removed by sterilisation but may also destroy any nutrients)(15), and high autofluorescence and intrinsic scattering which prevents study by many microscopical methods(16). These drawbacks can be circumvented using transparent refractive index-matched materials to create customisable soil scaffolds.

Transparent soil environments were primarily developed to study plant growth and plant-microbe interactions(17–19), requiring growth media that may have been suboptimal for either co-cultured organism. Moreover, most previous TS platforms have used either no refractive index matching solutions(20) or colloidal silica suspensions to refractive index match the soil substrate(18,19,21–23); however, these suspensions are often unstable over time, resulting in refractive index alterations owed to pH, salinity, and temperature changes.

We present the development of a chemically customisable, microscope-compatible, and environmentally analogous 3D culture method based on a Nafion™ scaffold, a fluorinated co-polymer of tetrafluoroethylene and perfluorosulfonic acid. We overcome the optical challenges of previous TS applications by using stable and tuneable sugar-based refractive index matching informed by the metabolic profiling of the test organism. Moreover, our workflow provides a flexible solution to facilitate the nutrient requirements of the test strain by permitting titration with various growth media. These two factors present our method as an adaptable technique for the study of microbes in a model 3D environment, especially by optical imaging.

## 5. Materials and Methods

The methods below outline our pipeline to construct a tailored 3D culture system for a given test microorganism. These steps involved the creation of a TS scaffold, the methods for which have been adapted from Downie *et al*.(18), and metabolic profiling to determine non-metabolisable sugars for stable refractive index matching.

The methods used to characterise TS in this work are not required to adapt the 3D culture method itself, and so are expanded in Supplementary Methods along with the imaging methods.

### 5.1 Production of a Nafion™ transparent soil scaffold for bespoke culture media

#### 5.1.1 Particle size reduction

A 10 g aliquot of Nafion™ NR-50 1100 EW (Ion Power GmbH, Germany) was cooled by submerging for 5 minutes in liquid nitrogen before being processed using a cryogenic grinder (6850, SPEX SamplePrep, USA) at the highest frequency (10.0 A.U.) for 2 minutes. Milled particles were sieved through a series of different pore sizes (500 μm, 850 μm, 1250 μm) (Fisher Scientific, UK) and particles larger than 1250 μm were re-processed as above until they passed through the 1250 μm sieve.

#### 5.1.1 Manipulation of Nafion™ surface chemistry to facilitate nutrient binding

The surface of milled Nafion™ particles was converted to an anionic form, therefore producing a hydrophilic surface and a negative charge to mimic the water retention profile of soil(18). This facilitated the downstream binding of nutrient cations. Milled Nafion™ was immersed in a solution of KOH (15% w/v), DMSO (35% v/v) and distilled deionised water (ddH_2_O) (R.2.0/200, Purite Ltd., UK) and incubated at 80°C for 5 hours. The conversion buffer was replaced with ddH2O and incubated at room temperature (RT) for 30 minutes. The particles were washed three times with ddH2O and incubated in 15% (v/v) HNO_3_ at RT for 1 hour. Particles were washed three times with ddH_2_O and incubated in 15% (v/v) HNO_3_ at RT overnight.

Inorganic and organic impurities were removed by washing in ddH_2_O three times and replacing with 1 M H_2_SO_4_. The suspension was incubated at 65°C for 1 hour before cooling down to RT. The H_2_SO_4_ was replaced with ddH_2_O and again incubated at 65°C for 1 hour before cooling to RT. Organic impurities were removed by washing three times with ddH_2_O and submerging in 3% (w/v) H_2_O_2_ and incubating at 65°C for 1 hour. The suspension was cooled to RT and washed three times with ddH_2_O and stored in ddH_2_O until required.

### 5.2 Tailoring transparent soil to produce a bespoke culture medium for *Streptomyces coelicolor*

Streptomycete culture was maintained by titration of TS with Supplemented Minimal Medium (SMM)(24) using L-arabinose, a common plant-derived sugar found in soil environments(25), used as the sole carbon source. Processed Nafion™ particles were titrated by submerging in fresh SMM and shaking for 30 minutes at 30°C, 225 rpm. The pH was measured after 30 minutes, and the spent medium was replaced with fresh SMM before again as above. This process was repeated until the pH of the spent medium was equal to that of fresh SMM broth (pH = 7.02). SO3H^+^ exchange sites were saturated after six titres and the pH stabilised. The particles were then refractive index matched before inoculation (Section 6.5).

### 5.2 Tailoring transparent soil to produce a bespoke culture medium for *Bacillus subtilis*

The above method of nutrient titration was adapted for *B. subtilis* culture using Spizizen Minimal (SM) medium(26) with D-glucose used as the sole carbon source. SO_3_H^+^ exchange sites were saturated typically after six cycles once the pH stabilised. The particles were then refractive index matched before inoculation.

### 5.2 Metabolic profiling to identify non-metabolisable sugars for refractive index matching

Logan & Berkley showed that *B. subtilis* is unable to metabolise certain carbon sources, but the specific strain studied was unclear(27). We performed a targeted metabolic screen to determine the carbon utilisation of *B. subtilis* 168 strains for refractive index matching candidates. Four sugars were selected from Logan and Berkely’s original screening, D-glucose, D-sorbitol, D-xylose and D-arabinose, and supplemented to SM medium (final [0.2% (w/v)]). A positive control of LB medium was used.

A 96-well optical bottom plate (265300, ThermoFisher Scientific, USA) was prepared with 198 μl of either SM medium supplemented with a test carbon source, or LB (with selective antibiotics where required). Overnight cultures *B. subtilis* 168 and JWV042 were grown in LB broth before diluting in fresh LB broth and growing to OD_600_ = 0.5. The cells were washed twice with sterile 1x SM buffer to remove any nutrient carryover. Two micro-litres of the washed cell suspensions were added to the respective wells before the plate was loaded into a Synergy™ HTX multi-mode plate reader (BioTek Instruments Inc., US) set to medium orbital shaking at 30°C while measuring OD_600_ every 15 minutes for 24 hours. Blanks containing sterile SM medium with each carbon source and sterile LB were also included. The specific growth rate (μ *h^-1^*) was calculated to quantify the growth on different nutrient sources and empirically identify non-metabolisable sugars, as described previously using Equation 1, where; *N* = the number of cells (OD_600_) at the final measurement timepoint (*t*) and *N_0_* = number of cells (OD_600_) at initial measurement timepoint (*t_0_*).

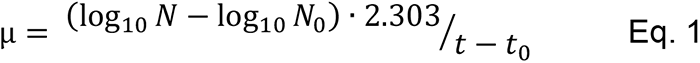

Although the method proposed was tailored to *B. Subtilis*, similar approach could be used to design the RI matching solution for other microorganisms.

*Streptomyces coelicolor* M145 is a well-documented in its inability to metabolise sucrose(28,29), which was verified by assessing the central carbon metabolism using the Kyoto Encyclopaedia of Genes and Genomes (30).

### 5.3 Refractive index measurements of Nafion and refractive index matching candidates

The refractive index of naïve Nafion™ particles was measured using Jamin-Lebedeff interferometry (Supplementary Methods), and the measured value was used as a target for candidate refractive index matching solutions. The refractive index of matching solutions was measured using an Abbe refractometer (Billingham & Stanley Ltd., UK) calibrated with methanol at 21°C (λ = 589 nm). A standard curve of concentration versus refractive index was prepared for each candidate in triplicate. Percoll^®^ was measured from 1% to 100% (v/v) diluted in distilled water (dH2O), sucrose was measured from 1% to 100% (w/v) diluted in SMM, D-xylose and D-arabinose were both measured from 1% to 50% (w/v) diluted in SM medium. Linear regression determined the concentration of refractive index matching agent to be supplemented into the final 3D growth medium to refractive index match Nafion™.

## 6. Results

### 6.1 Characterisation of Nafion™-based transparent soil for a customisable microbial culture platform

The optical properties of chemically processed Nafion^TM^ TS were verified before developing a bespoke culture platform (Figure 1). Freeze-fractured, surface-modified, and sulforhodamine-B-stained TS was imaged to verify the size/shape distribution of TS particles (Figure 1a). All particles measured 500 µm to 1250 µm in diameter, with irregularly shaped edges, as desired and in agreement with previous TS applications(18,19,21–23). The refractive index of processed TS particles was calculated using the phase information acquired with a Jamin-Lebedeff interferometer (Figure 1b, Supplementary Figure 1). The depth (*d*) of the measurement region (white cross in Figure 1b), chosen due to its relative flatness and therein more reliable measurement, was 210 µm. The phase retardation (Γ) at the measurement region was approx. 1400 nm. The refractive index was calculated using the above values and the known refractive index of the mounting medium, distilled water (*n_1_* = 1.333), resulting in the refractive index of Nafion^TM^ TS to be approx. 1.340, which concurs with measurements of other Nafion^TM^ polymers(31).

**Figure 1.**
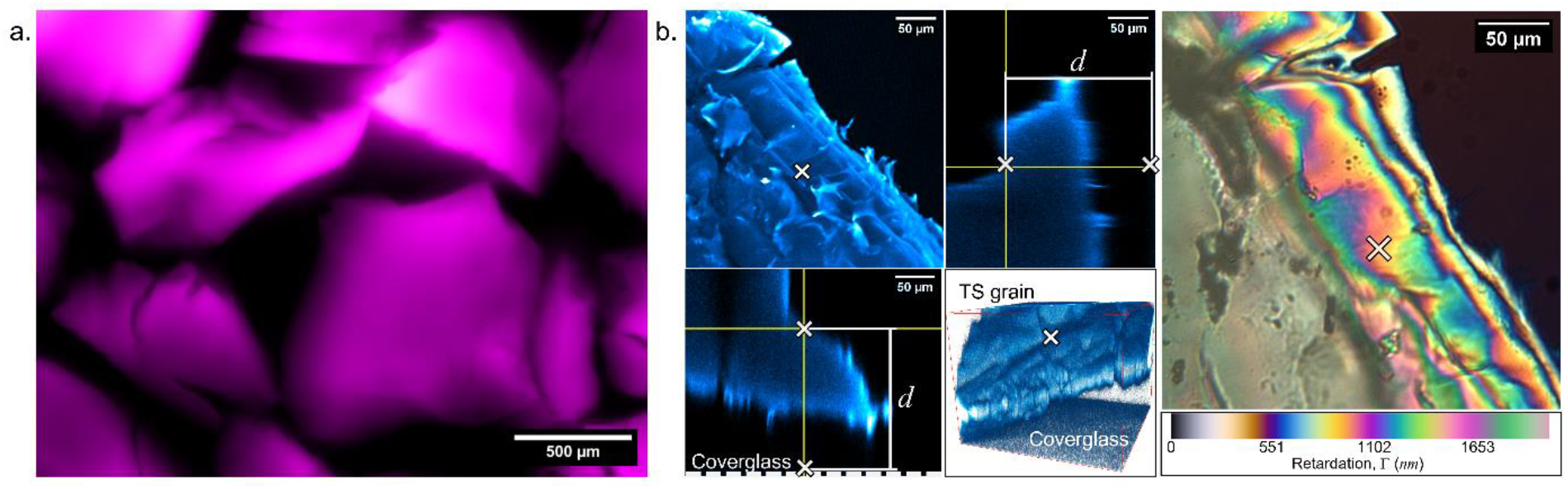
Optical characterisation of Nafion™-based transparent soil. **(a)** a widefield epifluorescence image showing a representative batch of cryomilled Nafion™ prior to chemical processing. The particles have been stained with sulforhodamine-B and false-coloured in magenta. **(b)** The refractive index of cryomilled Nafion™ was measured using Jamin-Lebedeff interferometry. A confocal *z*-stack maximum intensity projection, orthoview, and 3D reconstruction (cyan) provided the depth (*d*) of the measurement region (denoted by the white cross). The Jamin-Lebedeff image of the Nafion™ particle exhibited several interference orders which corresponded to the Michel-Lévy scale (bottom), phase retardation (Γ) = 1400 nm.

### 6.2 Exploiting non-metabolisable sugars for stable, inexpensive, and accessible refractive index matching

The conventional approach to refractive index matching in TS systems is by supplementation of colloidal silica. Commercial silica preparations can destabilise over long periods or from environmental changes, which can be caused by microbial metabolism. A more stable alternative was required for refractive index matching.

Carbon catabolism has been well studied in many bacterial species and provided a starting point to determine the optimal non-metabolisable sugar for TS refractive index matching. We compared their growth in LB medium to SM medium, with each sugar as a sole carbon source. The specific growth rate was calculated; *B. subtilis* 168 and JWV042 were unable to metabolise D-arabinose and D-xylose (Figure 2a, Supplementary Figure 2). The small growth rates observed were consistent with measurement errors over time compounded by the build-up of condensation on the lid of the plate.

Our other test organism, *S. coelicolor* M145 and its derivatives have a well-documented inability to degrade sucrose(28,29). Therefore, we reliably proceeded with sucrose to refractive index match any streptomycete experiments.

A standard curve of Percoll^®^, a commonly used refractive index-matching medium for TS, was set up to measure the refractive index using an Abbe refractometer (Figure 2b), and it was determined that a 35% (v/v) Percoll^®^ solution was required to match Nafion^TM^.

We determined the concentration of each sugar required to refractive-index match Nafion™ TS by measuring the refractive index of the three identified non-metabolisable sugars. Standard curves of sucrose, D-arabinose, and D-xylose (Figure 2c-e) revealed that 6.6% (w/v) sucrose was required to refractive index match TS for *S. coelicolor* observations and that 3.8% (w/v) D-arabinose or 5.1% (w/v) D-xylose was required for *B. subtilis* observations.

**Figure 2.**
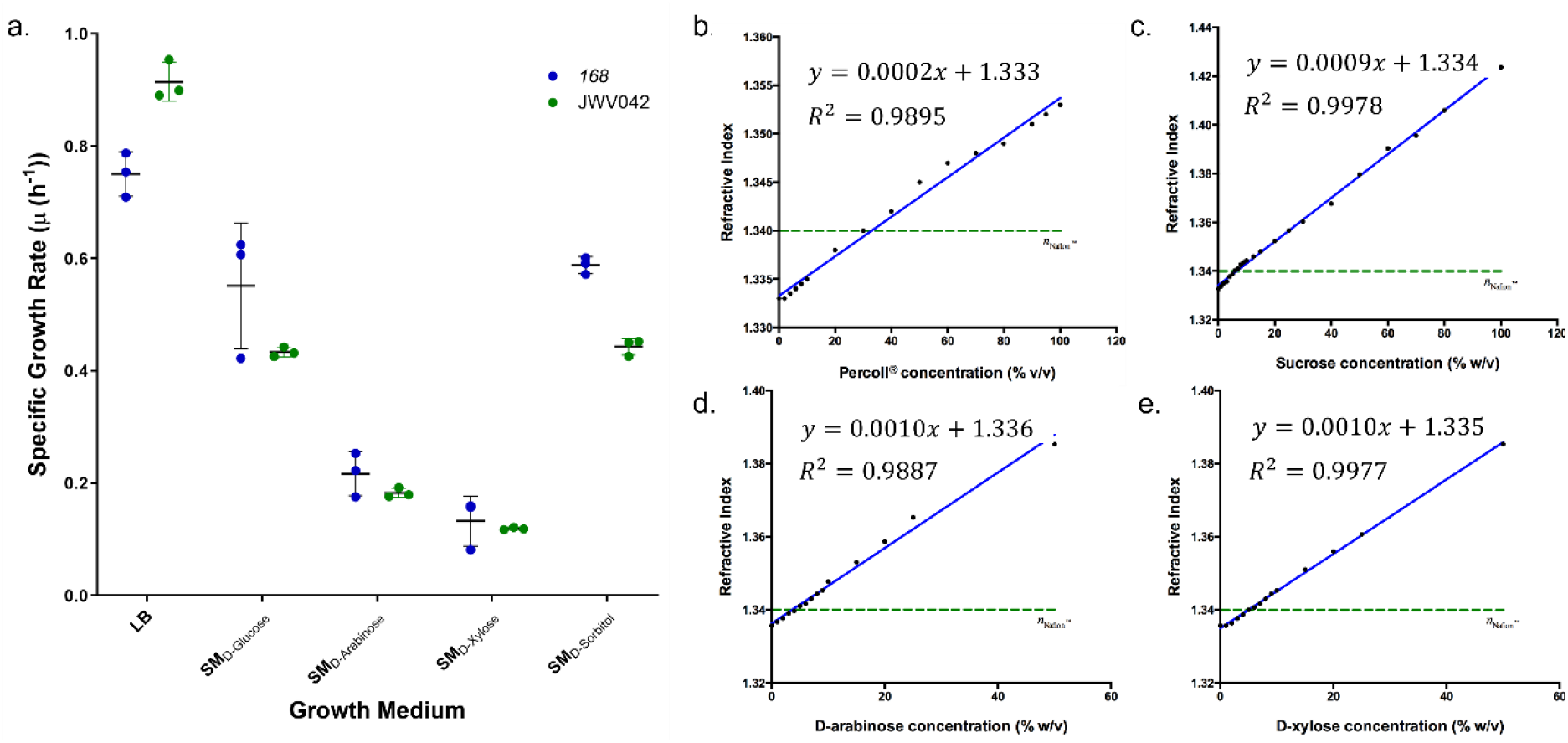
Identification of the optimal refractive index matching sugar by metabolic profiling. **(a)** The specific growth rate (*µ (h^-1^)*) of *B. subtilis* 168 and JWV042 in different minimal media containing different sole carbon sources compared with growth in LB. Strains in LB had significantly higher growth rates compared to incubation in minimal media. SM_D-arabinose_ or SM_D-xylose_ exhibited minimal growth, while SM_D-glucose_ or SM_D-sorbitol_ resulted in approximately 50% reduction in growth compared to LB. **(b)** A standard curve measuring the refractive index of Percoll^®^ dissolved in distilled water. The required concentration to refractive index match Nafion™ was **(b)** 35% (w/v) for Percoll^®^, 6% (w/v) for sucrose, **(d)** 3.8% (w/v) for D-arabinose, **(e)** 5.1% (w/v) for D-xylose. Blue fit = standard curve, green line = refractive index for Nafion™.

### 6.3 Implementation of bespoke 3D culture platform to image the colonisation behaviour of soil microbes

Following the identification of a suitable non-metabolisable sugar for refractive index matching, the 3D culture system could then be tailored for the study of individual strains of bacteria. We selected two phylogenetically distinct soil organisms to demonstrate the applications of this method, *B. subtilis* and *S. coelicolor*. A defined liquid growth medium was selected for each strain to maintain low levels of autofluorescence that would not prohibit downstream applications involving fluorescence. The nutrient-rich medium, Yeast Extract Malt Extract (YEME), was initially selected for streptomycete culture in transparent soil. However, YEME quickly turned transparent after the first addition and the Nafion™ particles were dyed yellow, indicating that medium components had saturated the surface of the soil particles.

As our endpoint involved imaging using the Mesolens(32), we first measured the autofluorescence intensity of Nafion™ TS using the Mesolens (Figure 3, Supplementary Figure 3), but also conducted measurements using a standard confocal laser scanning microscope (Supplementary Figure 4). Mesoscopic imaging revealed that chemically processed Nafion™ alone (“naïve”) had low basal autofluorescence, while TS imbibed with media containing yeast extract, YEME, exhibited 28-fold higher autofluorescence intensity. Autofluorescence intensity from SM_D-glucose_ was 43.23% lower than that of TS titrated with yeast extract-based media. Autofluorescence measurements of the same particle treatments using a routine confocal microscope revealed negligible autofluorescence signals for all treatments, meaning that more complex media could be explored for routine microscopy applications. The reusability of the 3D culture platform was tested by processing TS particles that had previously been stained with sulforhodamine-B. The dye was sufficiently removed following reprocessing of the stained Nafion™ and recycled TS had similar autofluorescence levels to naïve TS. Overall, results suggest that the use of media lacking yeast extracts is the best practice to minimise autofluorescence for downstream optical imaging applications and that our 3D culture platform can be repurposed for multiple experiments.

**Figure 3.**
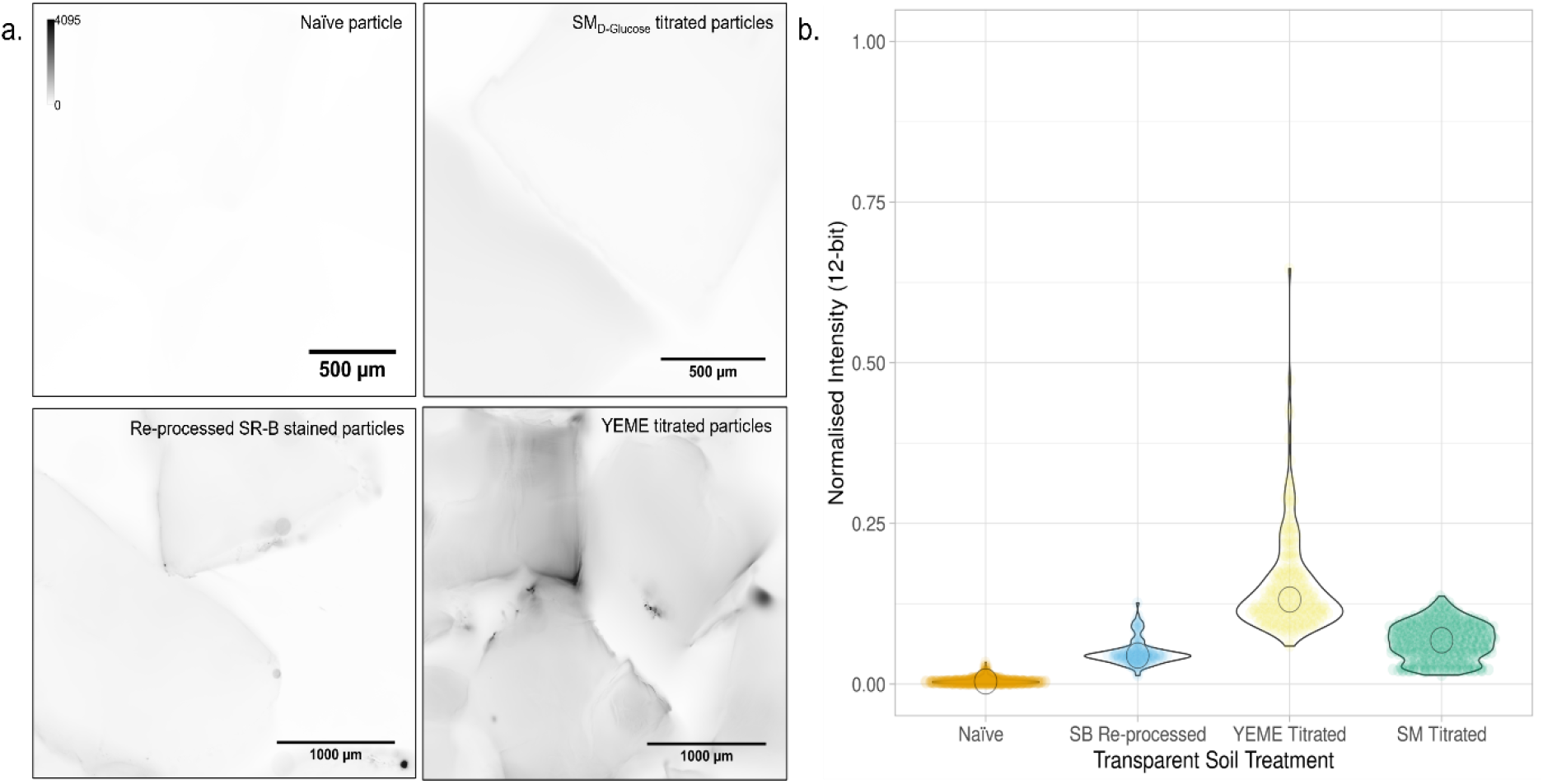
Autofluorescence of Nafion™-based TS depends on the components of the growth medium. **(a)** Mesolens images of chemically processed Nafion™ particles acquired using 490 nm excitation. Particles tested were naïve, recycled after being stained with sulforhodamine-B, titrated with SM_D-glucose_ medium, or titrated with YEME medium. The intensity scale was inverted to highlight autofluorescence for presentation. **(b)** Quantification of the autofluorescence signal from each TS specimen is shown in (a), where YEME-titrated TS had the highest autofluorescence intensity.

With an inert refractive index matching solution and the requirement for a defined liquid medium identified, SM-titrated TS matched with 3.8% D-arabinose was selected for *B. subtilis* imaging and SMM-titrated TS matched with 6.6% sucrose was used for *S. coelicolor* imaging. The application was demonstrated with cultures incubated in their respective tailored 3D culture medium and imaged using either a routine confocal laser scanning microscopy or the Mesolens setup in widefield epifluorescence or transmission brightfield mode. We demonstrated the application of our customisable 3D culture platform to study the colonisation behaviours of B. subtilis and S. coelicolor (Figure 4) in comparison to their routine growth behaviours under traditional laboratory culture condition (Supplementary Figure 5).

*Bacillus subtilis* exhibited two distinct colonisation behaviours when cultured transparent soil. Figure 5a shows a homogeneous but sparse colonisation of individual cells over the entire surface of TS grains in 3D and the presence of isolated biofilms in protected sites, such as grooves and cracks on the TS particle surface. *Streptomyces coelicolor* displayed one colonisation behaviour (Figure 4b). Dense microcolonies formed in protected regions, with hyphae extending from a dense mycelial cluster.

The prolonged viability of bacteria was tested by recovering *B. subtilis* JWV042 cells following seven days of incubation in a 3D culture scaffold. Cells were recovered and grown on rich medium and observed by widefield epifluorescence microscopy to ensure that cells remained viable during culture in the TS system (Supplementary Figure 6). Cells were easily recovered, regrown, and displayed the same phenotype as their initial inoculum, showing that our culture platform could maintain cell viability over multiple days.

**Figure 4.**
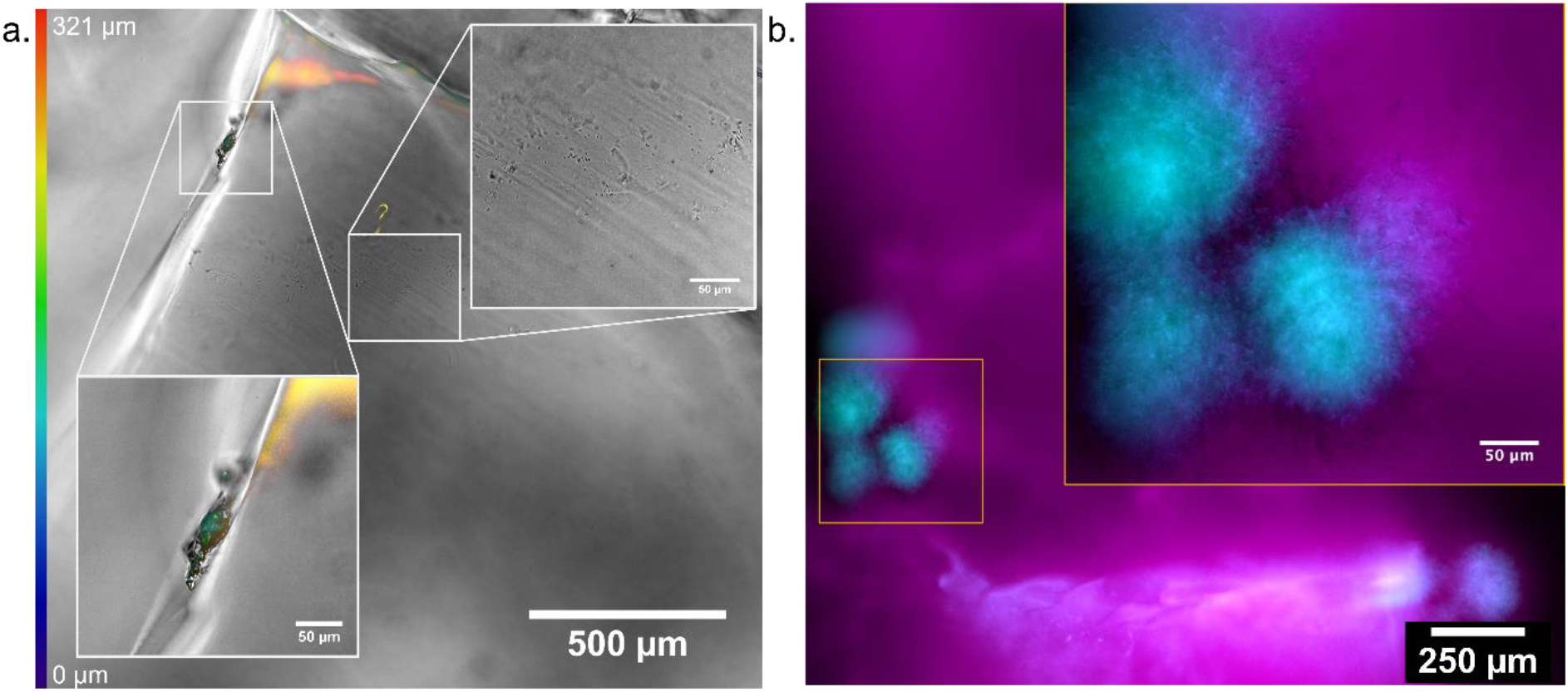
Growth behaviours of soil microbes in a transparent 3D culture environment. **(a)** A merged widefield epifluorescence and brightfield transmission Mesolens showing colonisation of *B. subtilis*. A *z*-coded colour table was used to show the 3D colonisation of the cells over the entire surface of the particles. Cells either grew as a uniform sparce covering or in discrete biofilms. **(b)** A widefield epifluorescence Mesolens image *S. coelicolor* grew as isolated microcolonies in protected regions in or between soil particles (*S. coelicolor idh*::*gfp* = cyan, TS autofluorescence = magenta).

## 7. Discussion

We report a methods pipeline to create a customisable, microscope-compatible, and environmentally imitative 3D culture platform for the growth and observation of soil microbes. Previous applications of transparent soil methods have been demonstrated in plant science(18,19,21,22) and interkingdom co-culture(20,23) but showed limitations; namely, difficulty in maintaining optical transparency, sub-optimal bacterial growth media, and understudies optical properties of TS. Our method optimises the transparency of the scaffold, manages autofluorescence, and provides scope for application to the culture of other soil microbes, allowing for *in situ* observation of colonisation dynamics and behaviours that may be impossible to replicate with current 2D laboratory culture methods.

We performed robust optical characterisation of our TS system and provide means to implement a custom refractive index matching process to render the substrate transparent. We demonstrated its use by culturing two phylogenetically distinct soil bacteria and present this as a flexible method for the soil microbiologist to adapt to use with their study organisms. Our pipeline is easily adaptable for the microbiologist to transfer to their test organism of choice and has the potential for future applications using fastidious isolates, investigating polymicrobial interactions, phenotyping of monocultured populations in representative environmental conditions, and mimicking extreme soil environments.

We first characterised the optical properties of Nafion™ transparent soil to inform the downstream chemical processing and refractive index matching requirements. It is important to note that optical characterisation is not routinely required for adapting our method by other users. Users may use our calculated values to optimise their own set-up. A refractive index of 1.340 for Nafion™ TS was calculated using Jamin-Lebedeff interferometry, agreeing with other measurements of bulk Nafion™(31). Jamin-Lebedeff interferometry was used as it provided enhanced sensitivity for refractive index measurements of soil substrates over, for example, Abbe refractometry which is typically used for liquid samples(33).

Nafion™ was selected due to its commercial availability, documented use for bioimaging, stability during autoclave sterilisation, low refractive index, potential for particle size refinement, and ion exchange capacity for nutrient imbibement(18,34). Using Nafion™ surmounted the optical and material complexities and costs of previously published artificial soil scaffolds, such as those using cryolite(20,35), fluorinated ethylene propylene(36), silica(37,38), and hydrogels(39,40). Moreover, we demonstrated that our platform was compatible with conventional fluorescence microscopy. We also demonstrated the application of the Mesolens with our culture method to take advantage of the high spatial resolution over a large field of view(32).

Following optical characterisation, we demonstrated a straightforward pipeline for stable sugar-based refractive index matching. This overcame the transiency of colloidal silica matching solutions in mutable environments and provided an accessible method for refractive index matching. Recently, transparent soil microcosms were developed to study soil microbes using cryolite and Nafion™ slurries(20). However, this application did not use optimal refractive index matching for high-quality optical imaging, negating the potential for longitudinal study or high-resolution optical imaging.

Commonly available sugars were prime candidates for refractive index matching agents due to their stability and high refractive index in solution. However, it was imperative that the study organism could not metabolise said sugar, otherwise, the refractive index of the culture medium would vary over the growth period. The use of sugars as refractive index matching agents provided the advantage of increased stability over time, less sensitivity to the media changing pH or temperature during incubation, and a higher degree of flexibility to refractive index tuning due to the inherently high refractive index of sugars. Moreover, the sugar concentrations we used were consistent with routine culture media recipes and low enough to not introduce osmotic pressures on growing cells(28,29). Furthermore, sugars represent a significant financial benefit as opposed to colloidal silica suspension; for example, the use of sucrose saw a cost reduction of 97% compared to using Percoll^®^ (£2.00 per litre 6.6% (w/v) sucrose versus £868.00 per litre 35% Percoll^®^, at the time of writing).

Autofluorescence was visualised using conventional confocal microscopy and mesoscopy. The detected autofluorescence was negligible using a routine fluorescence microscope that would be commonly available to other users. We then quantified the autofluorescence using the Mesolens to provide insight into the effects of different base nutrient solutions. The Mesolens was chosen for these measurements due to its 25-fold enhanced collection efficiency over a conventional optical microscope of the same magnification; this is owed to the unique ratio of magnification to numerical aperture, which also provides high spatial resolution throughout a large imaging volume(32). Our data showed that imbibing media containing yeast extract increased the detected levels of autofluorescence from the TS substrate. Therefore, to provide maximum fluorescence imaging quality, users are advised to use minimal media which avoid the use of yeast extract components. Minimal media recipes such as these have been suggested to be more akin to soil conditions, making the platform more representative of environmental conditions. Moreover, we demonstrated that used TS preparations can be reprocessed and recycled so that the substrate may be reused.

We applied our tailored culture medium to two distinct soil microbe monocultures and observed their colonisation behaviours using fluorescence microscopy and mesoscopy. We observed distinct modes of colonisation in both organisms which verify their previously described environmental life cycles and provide evidence of viability maintenance over 7 days of incubation in our 3D culture medium. Previous studies noted that the culturability of soil microbes could be improved by growing them in mimetic low nutrient structured environments(41). We have delivered a solution to facilitate these requirements and demonstrated its application. Our application overcomes the limitations of several other methods; the high autofluorescence and opacity of sterilised soil(42–44), the poor spatial resolution of correlative confocal/MALDI-MSI (Matrix-Assisted Laser Desorption/Ionisation-Mass Spectrometry Imaging)(45), and the non-representative spatial structure of fibreglass(46,47) or poly(ethene glycol)-based hydrogel scaffolds(48).

Adaption of our 3D culture method presents multiple new possibilities for the field of microbial ecology and microbial imaging. We have demonstrated the use of this platform with phylogenetically distinct soil bacteria and visualised the difference in growth behaviour using traditional culture methods. Our workflow allows users to customise their own 3D culture medium to suit their organism of study. We described how users may determine the optimum refractive index matching medium by conducting or using pre-determined screens to identify non-metabolisable sugars. The current system could be adapted to study the colonisation or growth behaviours of different microbes, to visualise the physiological effects caused by changing environmental conditions, or to determine the effects of chemical elicitation of new behaviours *in situ*. Moreover, the scope of the platform could be expanded to include titration with growth medium produced from soil extracts to generate an improved model environment, or to facilitate the growth of multiple isolates to study competition and growth dynamics.

## 9. Author statements

### 9.1 Acknowledgements

We wish to thank Dr Morgan Feeney (University of Strathclyde, UK) for informative discussions on growth measurements. We wish to thank Dr Brad Amos for assistance with the Jamin-Lebedeff interferometer, Miss Kay Polland for her assistance with figure preparation, and Dr Paul Herron (University of Strathclyde, UK) for use of his epifluorescence microscope.

We thank Dr Rut Carballido-Lopez (Université Paris-Saclay, France) for the gift of the *B. subtilis* strains.

The methods workflow presented in the graphical abstract was created with BioRender.com (Licence Number: SR25JKZ3PC). The violin plots created for Figure 3 were generated using PlotsOfData (developed by Joachim Goedhart, University of Amsterdam, Netherlands).

LMR and GM were funded by the Medical Research Council (MR/K015583/1) and The Leverhulme Trust. LXD was funded by the European Research Council (647857-SENSOILS). PAH was funded by BBSRC (BB/T001038/1 and BB/T004126/1) and the Royal Academy of Engineering Research Chair Scheme for long term personal research support (RCSRF2021\11\15).

### 9.3 Data Availability Statement

The authors confirm all supporting data and protocols have been provided within the article or through supplementary data files. The datasets generated during and/or analysed during the current study are available from the corresponding author upon reasonable request.

### 9.4 Author Contributions

LMR acquired and analysed the data and developed the method, with recommendations from LXD. LMR and LXD cryomilled and stained the Nafion substrate. LMR performed all downstream nutrient titrations and growth experiments. LMR, PAH, and GM designed the study. All Authors prepared and revised the manuscript.

## 10. Supplementary methods

The methods below outline the characterisation of Nafion™-based TS in this work but are not required to adapt the culture platform itself.

### 10.1 Bacterial strains and general maintenance

Two representative soil bacteria phyla were selected to characterise our optically transparent 3D culture system, Actinobacteria and Firmicutes, and well-studied species were selected from both phyla; *Streptomyces coelicolor* and *Bacillus subtilis*, respectively.

*Streptomyces coelicolor* (M145) cultures were maintained on solid Mannitol-Soya Flour (MS) medium (20 g/l mannitol, 20 g/l soya flour, 20 g/l agar, 1 l tap water) and liquid cultures were grown in 2x Yeast-Tryptone (2x YT) broth (16 g/l tryptone, 10 g/l yeast extract, 5 g/l NaCl, 1000 ml dH_2_O). Yeast Extract Malt Extract (YEME) medium (3 g/l yeast extract, 3 g/l malt extract, 5 g/l peptone, 10 g/l glucose, 1000 ml dH_2_O)(49) was used to demonstrate the autofluorescence of Nafion™ imbibed with yeast-based media. The GFP-expression strain, *S. coelicolor* M145 containing *idh*::*gfp* in its native location, integrated on the chromosome and was maintained by supplementing all media with 50 μg/ml apramycin. M145-*idh*-*gfp*(50) was selected as a test strain for this work as *gfp* was translationally fused to *idh*, which is a highly expressed primary metabolic gene encoding an isocitrate dehydrogenase found throughout the cytosol(51).

Streptomycete culture in TS was supported by titration with Supplemented Minimal Medium (SMM) (2 g/l casaminoacids, 5.73 g/l TES buffer [25 mM], 2 ml/l NH_2_PO_4_ + K_2_HPO_4_ buffer (50 mM of each) [1 mM], 1 ml/l 1M MgSO_4_·7H_2_O [5 mM], 2 g/l L-arabinose, 0.2 ml/l trace elements solution*, 1000 ml dH_2_O, pH 7.2, *trace elements solution: 0.1 g/l each of ZnSO_4_·7H_2_O, FeSO_4_·7H_2_O, MnCl_2_·4H_2_O, CaCl_2_·6H_2_O, NaCl).

*Bacillus subtilis* 168 and JWV042 cultures were maintained on solid Lysogeny (Lennox) Broth (LB) medium and grown in LB broth (10 g/l tryptone, 5 g/l yeast extract, 5 g/l NaCl, 1000 ml dH_2_O). The photoprotein encoding plasmid was maintained in JWV042 by supplementing the growth medium with 5 μg/ml chloramphenicol. JWV042 carried a plasmid which had a translation fusion of *gfp* and *hbs*, a gene encoding for the non-specific DNA-binding protein HBsu, which was under the control of an endogenous promotor (*cat amyE*::P*_hbs_*-*hbs*-*gfp*)(52).

Bacillus culture in TS was supported by titration with Spizizen Minimal (SM) Medium (2 g/l (NH_4_)_2_SO_4_, 14 g/l K_2_HPO_4_, 6 g/l KH_2_PO_4_, 1 g/l Na_3_-Citrate·2H_2_O, 200 mg/l MgSO_4_·7H_2_O, 500 mg/l tryptophan, 5 ml/l 50% (w/v) D-glucose, 1000 ml dH_2_O).

### 10.2 Fluorescent staining of *S. coelicolor* for routine laboratory growth observation

To visualise the structure of *S. coelicolor* grown under conventional laboratory conditions, specimens were stained using a method known as Schwedock staining(53). Briefly, this method uses two fluorescent dyes, fluorescein isothiocyanate-wheatgerm agglutinin (FITC-WGA) and propidium iodide (PI), to label *N*-acetylglucosamine residues in the peptidoglycan cell wall and nucleic acids, respectively. Cultures were grown on 22 mm x 22 mm type 1.5 coverglass (631-0125, VWR International Ltd., UK) sterilised by immersing in 100% ethanol before passing through a flame. Sterile coverglasses were inserted into solid MS medium at a 45° angle relative to the medium surface. A 5 μl of inoculum of a dilute M145 spore suspension was pipetted on the coverglass-medium interface at the acute angle and incubated at 30°C in darkened conditions for 72 hours (when the erection of aerial hyphae occurred). The coverglasses were aseptically removed from the agar and placed with the mycelium facing upwards abridge two sterile wooden toothpicks in a staining dish. The mycelium was fixed by pipetting 500 μl of ice-cold fixative (2.8% (v/v) formaldehyde (344198, Merck Millipore, USA), 0.0045% (v/v) glutaraldehyde (340855, Sigma Aldrich, USA), pH 7.0) onto the coverglass. The coverglasses were then washed twice with ice-cold phosphate-buffered saline (PBS) (BR0014, ThermoFisher Scientific, USA) before being allowed to air dry. The mycelia were then rehydrated by adding 500 μl of ice-cold PBS and incubating for 5 minutes at room temperature (RT). The PBS was then replaced with 500 μl of ice-cold Glucose:Tris:EDTA buffer supplemented with 2 mg/ml lysozyme (L4631, Sigma Aldrich, USA) and incubated for 1 minute at RT before being washed with ice-cold PBS. Finally, the mycelia were incubated with a blocking solution of 2% (w/v) bovine serum albumin (BSA) (A8531, Sigma Aldrich, USA) in PBS and incubated for 5 minutes at RT.

All subsequent staining steps were performed in darkened conditions. The blocking solution was replaced with 500 μl of an ice-cold staining solution containing 2 μg/ml FITC-WGA (W834, Invitrogen, USA) and 10 μg/ml PI (P1304MP, Invitrogen, USA) suspended in 2% (w/v) BSA in PBS. The coverglass was then incubated with the staining solution for 2 hours at RT. Following incubation, the first staining solution was removed and the coverglass was washed eight times with 500 μl of a second ice-cold staining solution containing only 10 μg/ml PI in PBS. Before imaging, 10 μl of 40% (v/v) glycerol was used to mount the stained mycelium.

### 10.3 Characterisation of the optical properties of transparent soil

#### 10.3.1 Sulphorhodamine-B staining of Nafion™ particles

Cryomilled and chemically processed Nafion™ particles were fluorescently stained to visualise the size and shape distribution of particles. 500 μl of 1 μg/ml (1.8 nM) sulphorhodamine-B (S1307, Invitrogen, USA) was added to a sample of milled and chemically processed Nafion™ particles. The dye solution was then removed and placed in a refractive index matching buffer before imaging.

#### 10.3.2 Jamin-Lebedeff interferometry

A Jamin-Lebedeff interferometer (Zeiss, Germany) (Supplementary Figure 1) was used to determine the refractive index of Nafion™ particles. The polarisation optics were aligned before each use to ensure a defined Newton’s Series was present at thin regions of the sample. A 10x interference objective, complimentary condenser apparatus, and a C-mounted CMOS colour camera (DKF 33UX264, The Imaging Source, Europe GmbH, Germany) were used to acquire an interference imaging of the Nafion™ particles. White light illumination was sourced from a tungsten lamp. Single grains of cryomilled Nafion™ were mounted between two Type 1.5 coverglasses in dH_2_O. A region showing a clear separation of Newton’s interference orders was selected, and the colour and order of the measurement region were noted and compared to a Michel-Lévy chart to obtain the phase retardation.

To determine the depth of the sample at the measurement region, the specimen was carefully removed and inverted before placing it on an Olympus IX81 inverted microscope coupled to a FluoView FV1000 confocal scanning unit (Olympus, Japan). The particle autofluorescence was used to gain an overview of the surface in three dimensions. Images were acquired using a 10x 0.4 NA UPLXAPO objective lens (Olympus, Japan) and fluorescence emission was detected using a photomultiplier tube with spectral detection set to 525 nm with a 50 nm bandwidth. A 488 nm line from an Argon laser (GLG3135; Showa Optronics, Japan) was used as an excitation source. A 3D *z*-stack of the same area imaged using the Jamin-Lebedeff interferometer was acquired at Nyquist rate (Δ*z* = 1.50 µm). The upper and lower limits were used to measure the depth of the measurement region.

The refractive index of Nafion™ (*n_2_*) was calculated using the established relationship between the thickness of the specimen (*d*), the phase retardation (Γ) at the measurement region, and the known refractive index of the mounting medium (dH_2_O) (*n_1_*); *n*_2_ = ^Γ^⁄_*d*_ + *n*_1_ (54). For accurate measurement, the mounting medium should be as close to the specimen as possible (i.e., *n*_1_ = *n*_2_ ± 0.05), and so some optimisation is required for specimens of a completely unknown refractive index.

### 10.4 Imaging conditions

#### 10.4.1 Widefield epifluorescence microscopy

Conventional widefield epifluorescence and phase contrast microscopy were used to compare the cellular morphology and GFP expression before and after the growth of bacteria in a transparent soil environment. Images were acquired using an Eclipse TE-2000-S microscope with a 100x/1.30 NA CFI PLAN FLUO DLL Oil objective lens (Nikon, Japan) coupled to a digital CCD camera (C4742, Hamamatsu, Japan). Illumination for GFP excitation was sourced from a mercury arc lamp which was coupled into the epi-port of the microscope. A 530 nm ± 35 nm emission filter was used for fluorescence imaging (1CH81700, Chroma Technology Corporation, USA). Illumination for phase contrast microscopy was provided by a tungsten bulb light source.

#### 10.4.2 Confocal laser scanning microscopy

A confocal laser scanning microscope was used to verify that no reflection signal was detected from soil particles following refractive index matching and to assess the autofluorescence profile of the transparent soils. Particles were imaged using a Leica TCS-SP5-II confocal laser scanning microscope with a 5x/0.15 NA PL FLUOTAR objective lens (Leica, Germany). Incident light for reflection, autofluorescence excitation and transmission was provided by a 488 nm line from a Kr/Ar laser source (Coherent, USA). Reflection imaging was carried out by placing an 80/20 beam splitter in the detection path and restricting the PMT detection to 488 nm ± 5 nm. The autofluorescence signal was acquired by extending the same PMT detection from 520 nm to 620 nm. Transmission images were acquired by using the TCS-SPT-II in transmission mode and detecting transmitted light using a PMT detector

#### 10.4.3 Widefield mesoscopy

Brightfield transmission mesoscopy was achieved using a tungsten bulb light source and the condenser position, condenser diaphragm, and field iris set for Kohler illumination. Fluorescence excitation was achieved using a 435 nm or 490 nm LED from a pE-4000 LED illuminator (CoolLED, U.K.). A triple bandpass filter which transmitted light at 470 ± 10 nm, 540 ± 10 nm, and 645 ± 50 nm was placed in the detection pathway. The emission signal was detected using a VNP-29MC CCD camera with chip-shifting modality (Vieworks, South Korea) to capture the full FOV of the Mesolens at high resolution. Widefield mesoscopic imaging was carried out using water immersion (*n* = 1.33) with Mesolens correction collars set accordingly to minimise spherical aberration through refractive index mismatch.

### 10.5 Image analysis

The autofluorescence of Nafion™ particles following different surface treatments was analysed by first acquiring images of soil particles by widefield epifluorescence mesoscopy. As each test strain of bacteria encoded GFP, the autofluorescence intensity was measured in all samples by exciting with 490 nm LED at moderate power and exposure times (35% LED power, 500 ms exposure time). Autofluorescence intensity was quantified by first drawing multiple ROIs randomly over the surface of transparent soil particles. The mean intensity in each ROI was measured and normalised against the dynamic range of the camera sensor (0 – 4095 AU). The mean values for each surface treatment were then compiled in a violin plot using Plots of Data. Image analysis was performed using FIJI(55). Figures presented were linearly contrast adjusted for presentation purposes where required using FIJI.

### 10.6 Validating specimen viability in transparent soil media

A phenotypic screening method was used to determine if bacteria remained viable throughout incubation and growth in a transparent soil environment. Following growth and imaging of JWV042 in transparent soil, a 10 μl inoculum was taken from the soil culture and used to inoculate a 5 ml volume of sterile LB broth supplemented with 5 μg/ml chloramphenicol. The culture was then grown overnight at 37°C shaking at 225 rpm before being serial diluted in sterile LB broth to 1x10^-8^ and plated on solid LB medium supplemented with 5 μg/ml chloramphenicol and incubated for 18 hours at 37°C in darkened conditions. A sample was also taken from the overnight liquid culture for visualisation under a conventional widefield epifluorescence microscope to establish if cells recovered from transparent soil remained fluorescent. These cells were compared to cells which had been grown for 48 hours in liquid SM_L-arabinose_ medium. The cellular morphology was compared using the microscopy data, while colonial morphology was compared using the colonies on the serial dilution plates.

## 11 Supplementary Figures

**Supplementary Figure 1.**
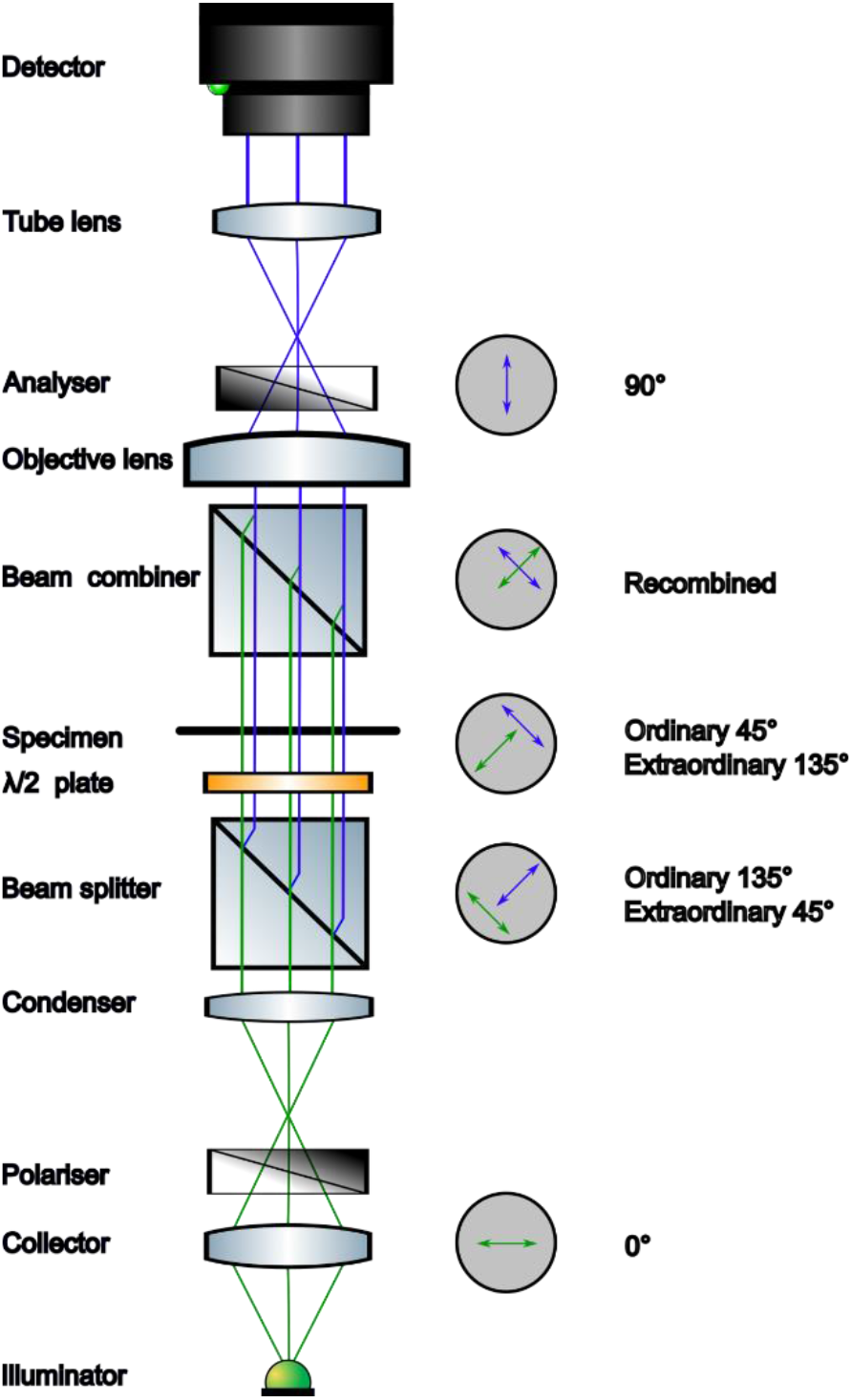
Schematic of a Jamin-Lebedeff interferometer. White light illumination is collected and transmitted through a polariser aligned parallel to the optical axis, before being transmitted through a beam splitter which splits the incident light into two paraxial rays: the measurement beam (blue) and the reference beam (green). A rotating half-wavelength plate is incorporated into the condenser apparatus, azimuthally rotating will control the alignment of the Newton’s series in the acquired interference image. Light passes through the sample, and the measurement and reference beams will undergo a relative phase retardation depending on their transmittance through a sample of depth, *d*. The two beams are re-combined by a beam combiner, which rotates azimuthally together with the objective to match the diagonal position of the crossed polariser and analyser. The objective lens then collects the re-combined light to form an image displaying the phase retardation as a coloured Newton’s series, as in a Michel-Lévy chart. The analyser is oriented to match the polariser and aligns the plane and axis of the combined beams to cause destructive and constructive interference between the two wavefronts. The light then passes through a tube lens prior to being detected, either by eye or by CCD or CMOS camera.

**Supplementary Figure 2.**
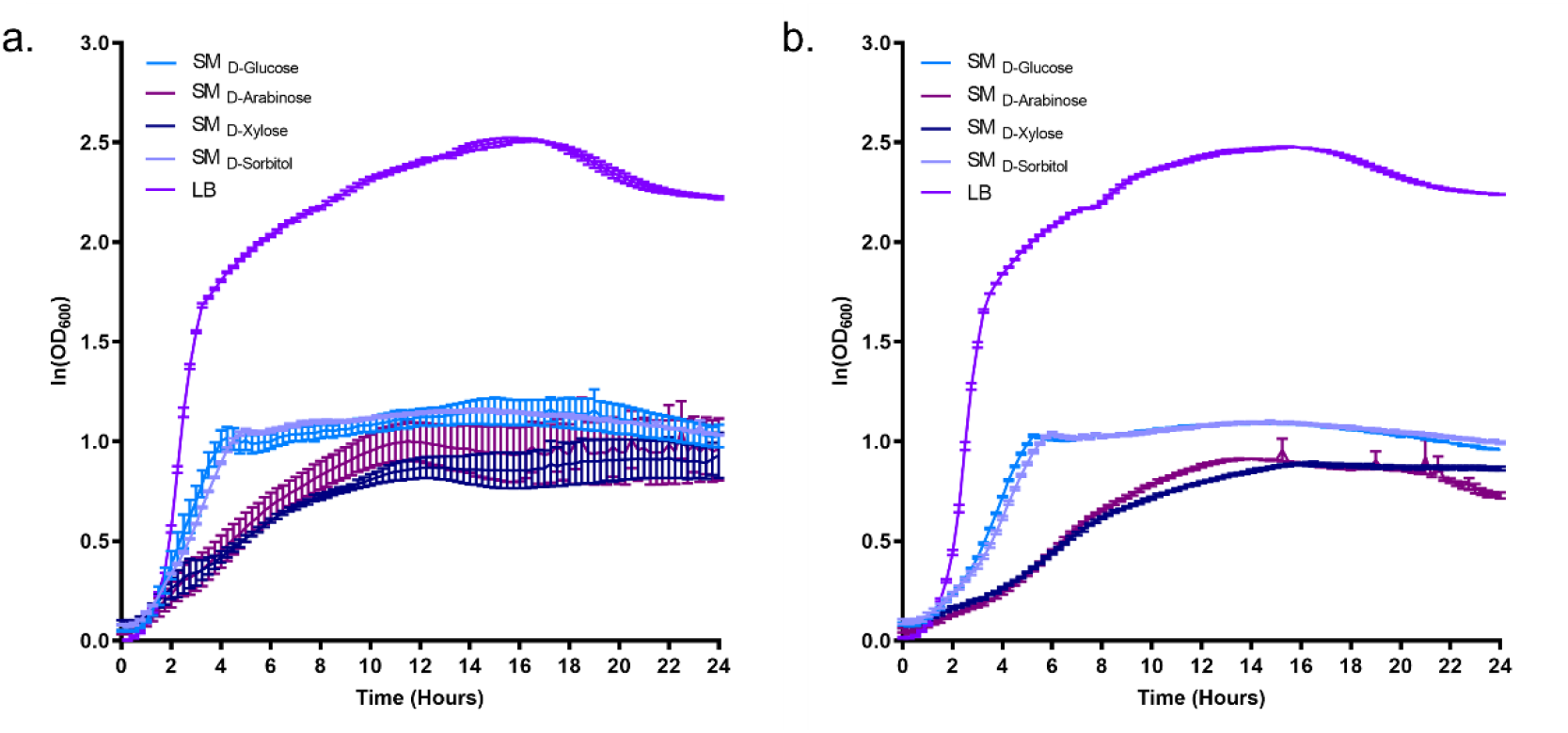
Growth curves of *B. subtilis* to determine non-metabolisable sugars. **(a)** A semi-log growth curve for *B. subtilis* 168 grown on rich medium (LB) and several SM media containing a single sugar carbon source. **(b)** A semi-log growth curve for *B. subtilis* JWV042 grown on rich medium (LB) and several SM media containing a single sugar carbon source.

**Supplementary Figure 3.**
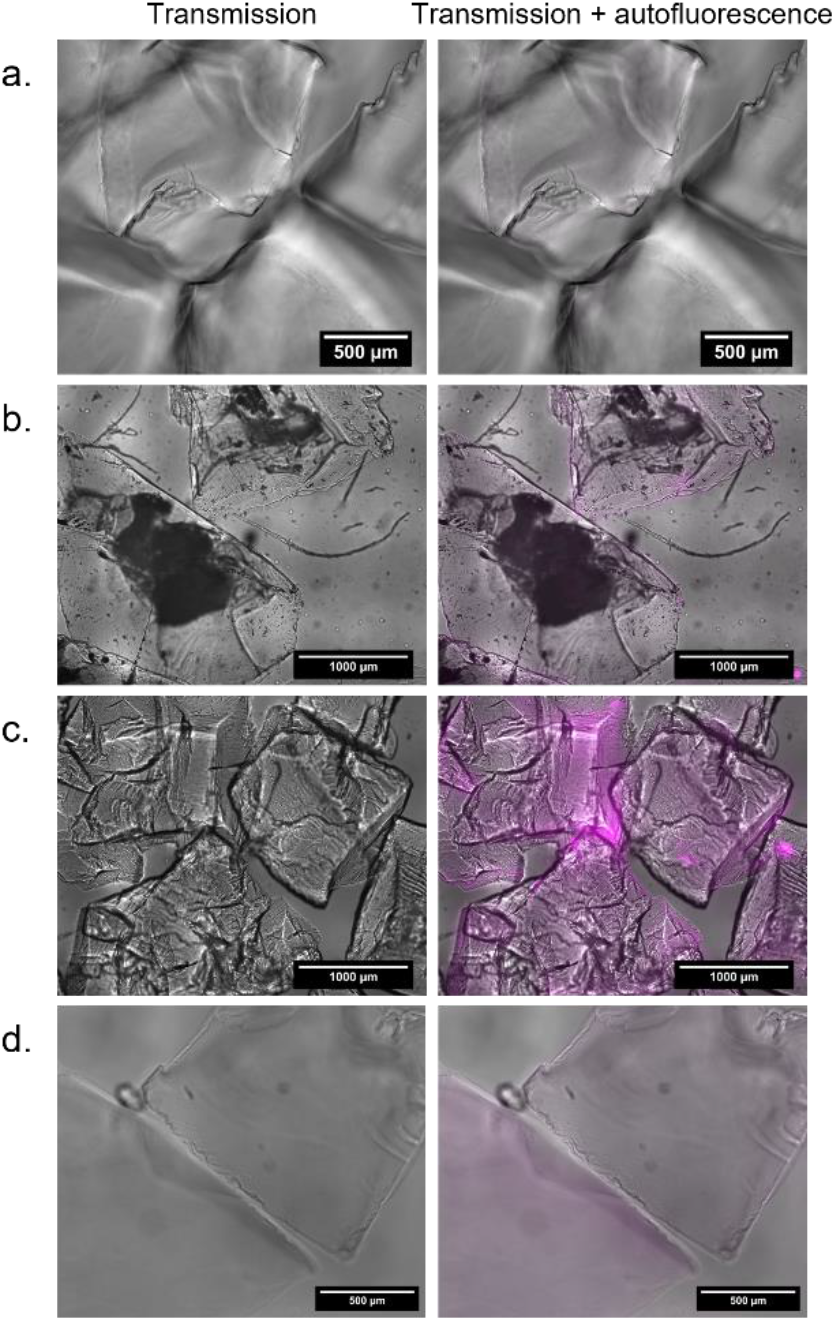
Merged brightfield and fluorescence images of TS autofluorescence acquired using the Mesolens. **(a)** Brightfield transmission (left) and merged fluorescence (right) images of naïve Nafion™-based TS. **(b)** Brightfield transmission and merged fluorescence images of recycled sulphorhodamine-stained TS. **(c)** Brightfield transmission and merged fluorescence images of YEME-titrated TS. **(d)** Brightfield transmission and merged fluorescence images of SM_D-glucose_-titrated TS. Brightfield transmission is presented in grayscale, autofluorescence is false coloured in magenta. Fluorescence images alone were used to quantify autofluorescence signals in Figure 3.

**Supplementary Figure 4.**
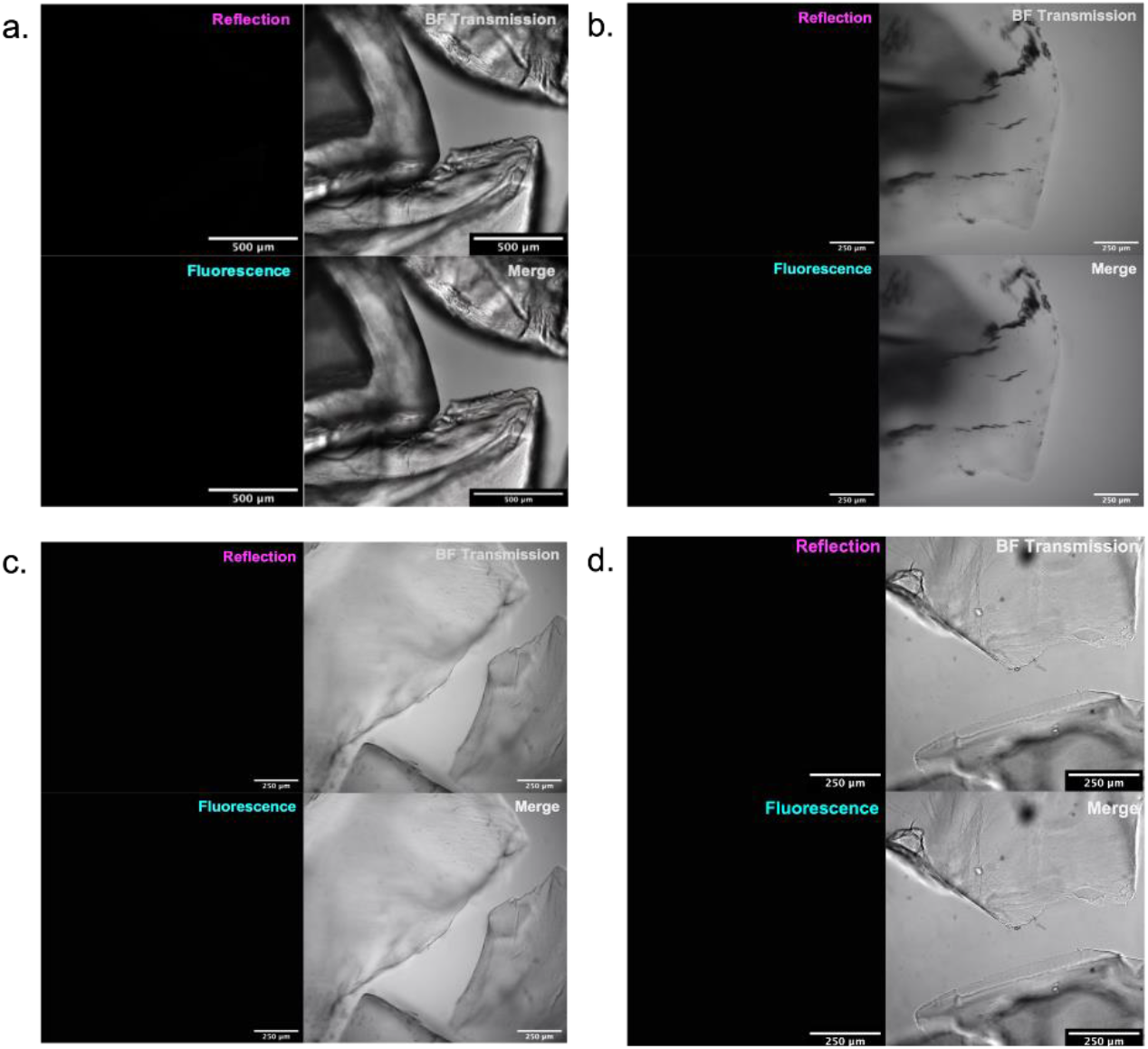
Autofluorescence of Nafion™-based TS acquired using a conventional confocal laser scanning microscope. Images acquired using a routine confocal microscope show negligible autofluorescence signal is detected in naïve **(a)**, recycled **(b)**, YEME-titrated **(c)**, or SM-titrated Nafion™ TS **(d)**. Brightfield transmission is presented in grayscale, autofluorescence is false coloured in cyan, and reflection is false coloured in magenta.

**Supplementary Figure 5.**
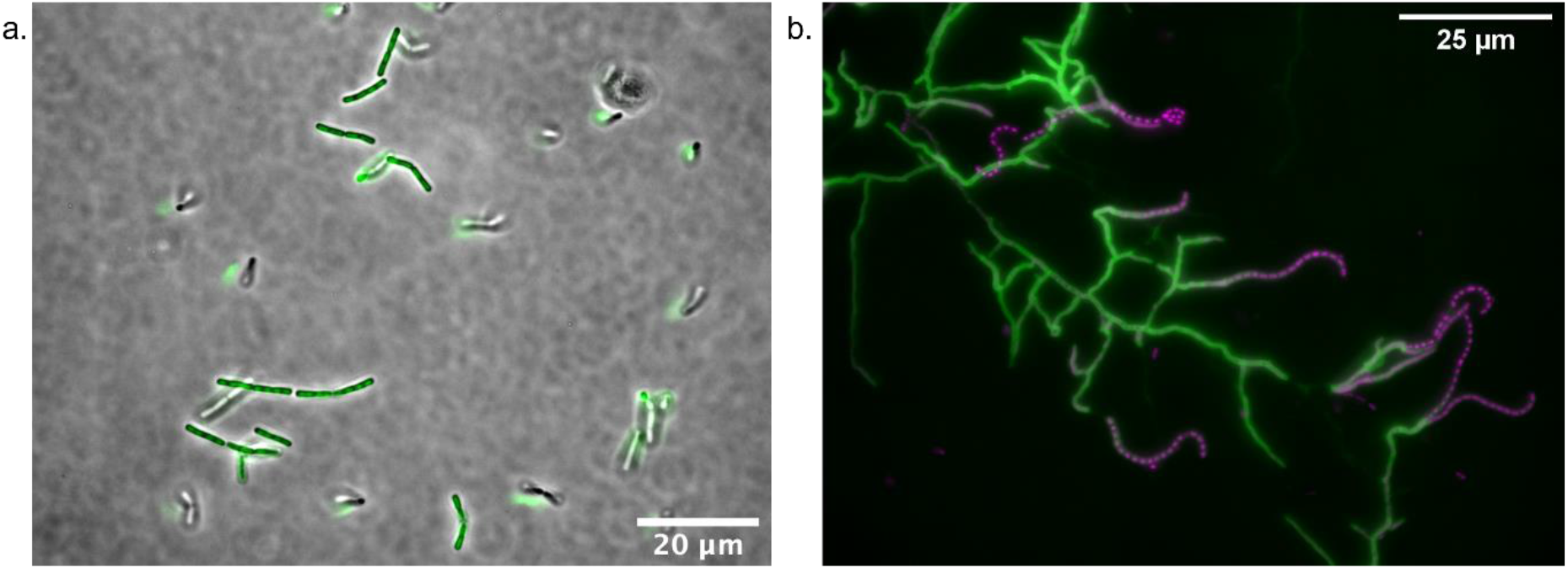
Normal laboratory growth behaviours of *B. subtilis* and *S. coelicolor* under traditional culture conditions. **(a)** *B. subtilis* JWV042 is routinely culture as large multimillimetre colonial biofilms on solid nutrient rich agar or, as shown here, as single cells or short chains in shaking liquid culture. Image shows phase contrast (grey) and Hbs-GFP fluorescence (green) **(b)** *S. coelicolor* routinely grows as a differentiated mycelium, with vegetative hyphae undergoing a series of regulated lifecycle changes to erect aerial hyphae and forming spore chains under nutrient-deprived conditions. Nucleoids are stained with propidium iodide (magenta) and peptidoglycan cell walls are stained with FITC-wheatgerm agglutinin (green).

**Supplementary Figure 6.**
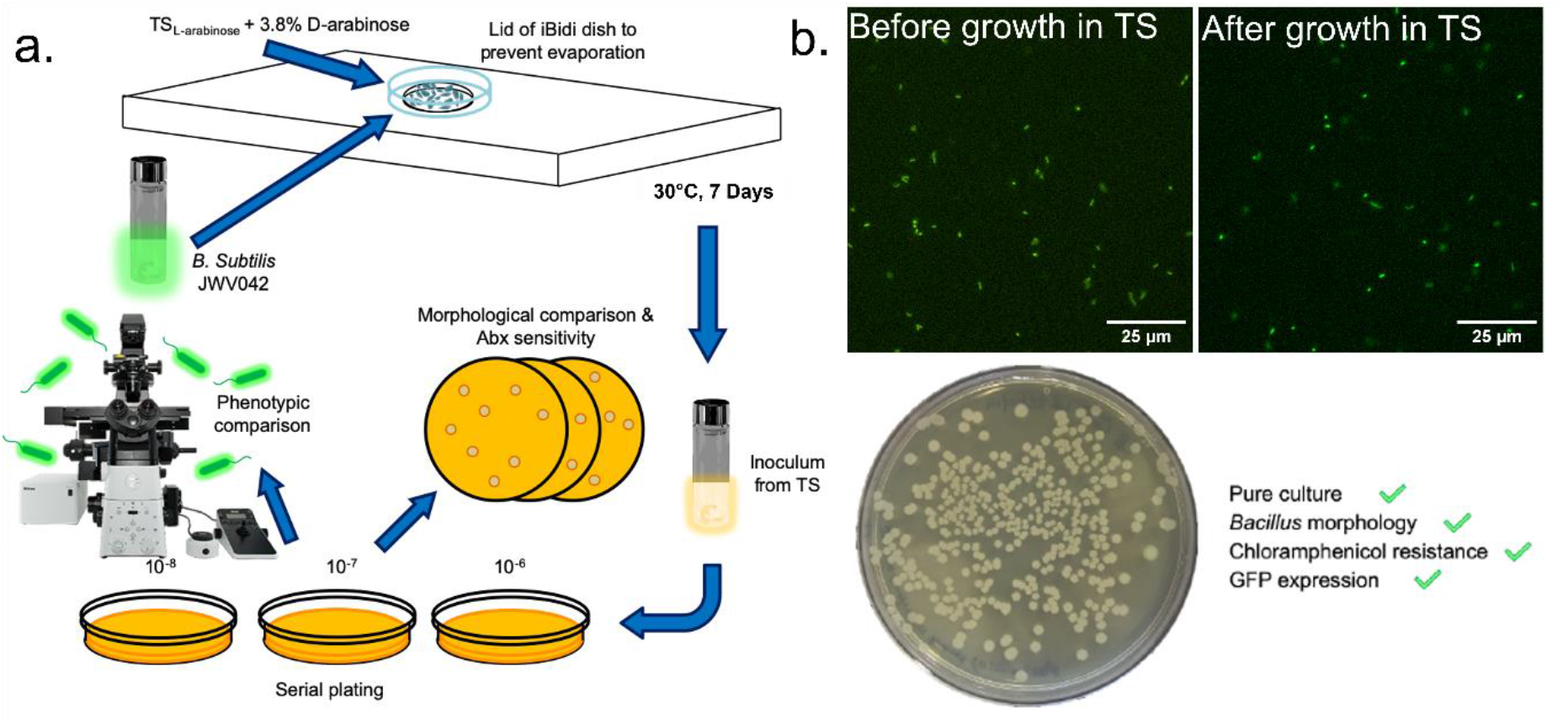
Cells remain viable and recoverable after prolonged periods using the tuneable TS culture platform. **(a)** A schematic of the recovery process for *B. subtilis* JWV042 after 7 days of culture in a custom 3D culture medium. Cells were growth for 7 days prior to removal and being serial plated on routine solid culture media. Recovered were grown in the presence of a selective antibiotic and imaged by widefield epifluorescence microscopy to confirm they had the same phenotype as their progenitor inoculum. **(b)** Comparative fluorescence micrographs showing the progenitor and recovered cells after 7 days in 3D culture, Hbs-GFP fluorescence is presented in green. Recovered cell phenotypes were maintained, as shown by plate-based observations.

